# Connectivity allometry is a robust organizing principle of the human functional connectome

**DOI:** 10.64898/2026.09.14.751437

**Authors:** Changwen Wu, Xiaofan Qiu, Junle Li, Suhui Jin, Xueyan Jiang, Jinhui Wang

## Abstract

A defining feature of the human brain is its multiscale organization, in which regional processes are integrated into coherent whole-brain network architecture. How local functional organization adapts to variation in this global architecture, however, remains largely unknown. Here, we addressed this question by applying an allometric scaling framework to resting-state functional magnetic resonance imaging data from four large-scale datasets spanning different cultures, cognitive states, developmental stages, and clinical conditions. We found that human functional networks were organized according to a robust and spatially heterogeneous connectivity allometry, whereby regional functional connectivity scales nonlinearly with whole-brain connectivity strength. This scaling architecture closely followed the cortical sensorimotor–association hierarchy, with lower-order systems exhibiting supralinear scaling and higher-order systems exhibiting sublinear scaling. Connectivity allometry was further associated with multiscale neurobiological organization, including regional metabolic properties, neurotransmitter receptor distributions, and transcriptomic landscapes, indicating that local–global functional scaling is embedded within fundamental biological gradients of cortical organization. Moreover, the scaling pattern and its biological associations were highly reproducible across independent cross-cultural cohorts. Importantly, connectivity allometry remained remarkably stable across working memory conditions regardless of cognitive load, emerged by late childhood, and was broadly preserved in children and adolescents with autism spectrum disorder despite region-specific alterations. Together, these findings establish connectivity allometry as a previously unrecognized organizational principle of the human functional connectome and provide a quantitative framework linking regional functional organization to whole-brain network architecture in health and disease.

## Introduction

Biological systems are organized across multiple spatial scales, raising a fundamental question of how local components adapt to variation in system-level organization. Rather than changing proportionally with global variation, individual components often exhibit nonlinear scaling relationships, a phenomenon known as allometric scaling [1,2]. In the human brain, allometric scaling has been extensively characterized for regional morphology, which have consistently been shown to scale nonlinearly with global properties (e.g., overall brain size) [3–8]. These findings indicate that the human brain follows robust local–global scaling laws at the regional level, providing crucial insights into organizational and adaptive principles underlying brain development and evolution [9,10]. However, whether these scaling principles extend beyond regional brain properties to the organization of large-scale brain networks remains largely unknown.

Human brain networks can be mapped in vivo using resting-state functional magnetic resonance imaging (R-fMRI), which enables quantifying functional connectivity (FC) of spontaneous neural activity among distributed brain regions [11] and has become one of the most widely used approaches for investigating large-scale brain organization [11,12]. Similar to regional morphology, FC exhibits substantial interindividual variability and systematic changes across the lifespan [13–15]. For example, a recent large-scale normative study has shown that global and regional FC follow coordinated yet distinct developmental trajectories. Moreover, accumulating evidence indicates that functional subnetworks mature asynchronously, with primary sensorimotor systems reaching mature configurations earlier than higher-order association networks [15,16]. These observations collectively suggest that local and global functional organization are tightly coupled but not simply proportional. Instead, different regions and cortical systems may contribute differently to variation in whole-brain connectivity, implying that regional FC may obey distinct allometric scaling relationships. Despite these observations, a quantitative framework for characterizing local–global scaling in functional brain networks is currently lacking.

In this study, we tested the hypothesis that human functional networks are governed by a spatially heterogeneous local–global scaling law, whereby regional FC strength scales allometrically with whole-brain global FC strength. To test this hypothesis, we analyzed R-fMRI data from the Human Connectome Project Young Adult (HCP-YA) cohort and independently validated our findings in the Southwest University Longitudinal Imaging Multimodal (SLEM) dataset. Using log–log regression, we estimated region-specific scaling coefficients to characterize the cortical topography of connectivity allometry. We further investigated how this scaling architecture relates to macroscale cortical hierarchy and microscale cortical organization, including neurotransmitter receptor distributions and transcriptomic landscapes. Finally, we examined the generalizability of connectivity allometry across cognitive states using task fMRI and assessed it alteration in autism spectrum disorder (ASD). Together, this work introduces a quantitative framework for characterizing local–global scaling in functional brain networks and establishes connectivity allometry as a previously unrecognized organizational principle of the human functional connectome. By linking regional functional organization to whole-brain network architecture across multiple spatial, biological, and behavioral scales, our framework provides a new systems-level perspective for understanding the principles governing human brain organization in both health and disease and establish a normative reference for identifying abnormal network reorganization associated with brain development, aging, and neurological or psychiatric disorders.

## Results

### Overview of the allometric scaling framework in the human functional connectome

In this study, we investigated whether regional FC follows a systematic allometric scaling rule relative to whole-brain global connectivity. To test this, we modeled the relationship between regional FC strength (*S_nodal_*) and whole-brain global FC strength (*S_global_*) using a log–log regression framework, from which we derived a region-specific scaling coefficient, *β*_1_. Within this framework, *β*_1_ = 1 indicates isometric scaling, such that regional connectivity changes proportionally to global fluctuations. By contrast, deviations from unity indicate allometric scaling: *β*_1_ > 1 (positive allometry) reflects a disproportionately faster change in regional connectivity relative to global connectivity, whereas *β*_1_ < 1 (negative allometry) indicates a slower rate of change in regional than global connectivity.

We evaluated this framework across four complementary datasets by constructing individual functional brain networks with 400 cortical regions. First, we used the HCP-YA resting-state dataset (Western; N = 413) as the discovery cohort for mapping the cortical topography of connectivity allometry. To move beyond descriptive characterization, we examined the resulting allometry map in relation to both macroscale and microscale cortical organizing principles, including cortical hierarchy, metabolic architecture, neurotransmitter distributions, transcriptomic profiles, and meta-analytic activation maps. We then independently replicated the full analytical pipeline in the SLEM dataset (Eastern; N = 464) to test cross-cultural generalizability. Finally, to demonstrate the broader implications of FC allometry, we examined whether and how it is modulated across cognitive states using HCP-YA task dataset (N = 400) and whether it is altered in a clinical condition using the Autism Brain Imaging Data Exchange (ABIDE) dataset (N = 856) in autism spectrum disorder (ASD). Together, this multi-dataset framework establishes connectivity allometry as a robust and fundamental organizing principle of the human functional connectome.

### Cortical topography of FC allometry

As shown in Figure 1A, FC allometry showed a spatially heterogeneous cortical distribution. This topography was highly consistent between the Western HCP-YA cohort and the Eastern SLEM cohort (*rho* = 0.832, *p*_spin_ < 0.001, Figure 1B), and showed striking inter-hemispheric symmetry (HCP-YA: *rho* = 0.644, *p*_spin_ < 0.001; SLEM: *rho* = 0.599, *p*_spin_ < 0.001; Figure S1) and high similarity between males and females (HCP-YA: *rho* = 0.857, *p*_spin_ < 0.001; SLEM: *rho* = 0.824, *p*_spin_ < 0.001; Figure S2). Testing for deviation from isometric scaling (i.e., scaling coefficient = 1) identified a specific set of regions showing significant allometric scaling between *S*_nodal_ and *S*_global_ [HCP-YA: 318 regions; SLEM: 247 regions; *p*_spin_ < 0.05, false discovery rate (FDR) corrected; Figure 1C)]. The allometric regions showed high spatial overlap between the two cohorts (Dice coefficient = 0.805; Figure 1D). Specifically, positive allometry was observed in 114 regions in the HCP-YA dataset and 88 regions in the SLEM dataset, primarily involving the precentral and postcentral gyri, paracentral lobule, and several occipital regions. Negative allometry was observed in 204 regions in the HCP-YA dataset and 159 regions in the SLEM dataset, predominantly involving the dorsolateral and medial prefrontal cortex, lateral temporal cortex, and inferior and medial parietal cortex.

**Figure 1.**
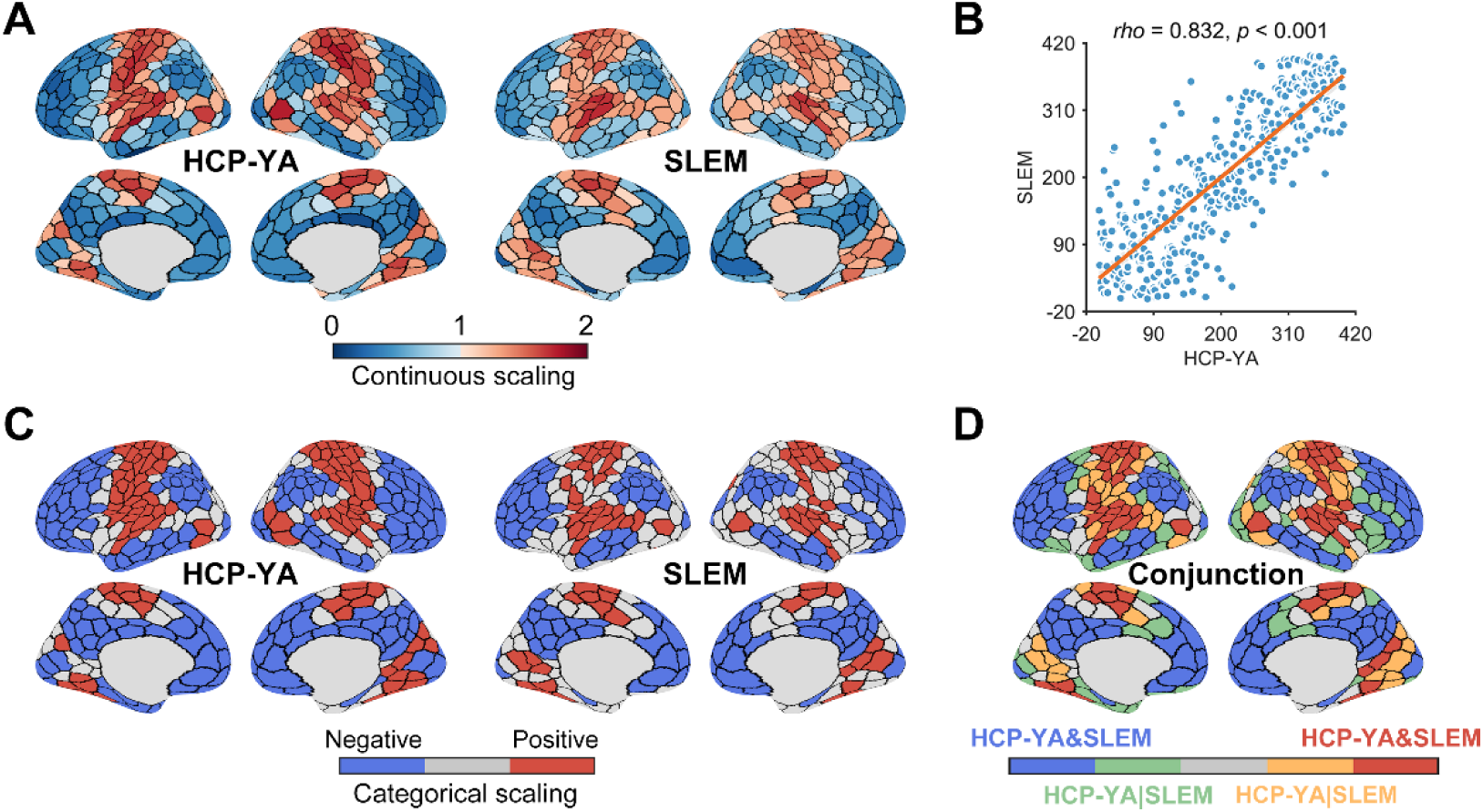
Cortical topography of FC allometry and cross-cohort consistency. **A**) Regional scaling coefficients showed a spatially heterogeneous distribution across the cortex in the HCP-YA and SLEM cohorts. **B**) A high positive correlation was observed between FC allometry maps derived from the HCP-YA and SLEM cohorts. **C**) A specific set of regions showed significant deviations from isometric scaling in the HCP-YA and SLEM cohorts. **D)** Regions of significant allometric scaling showed high spatial overlap between the HCP-YA and SLEM cohorts. HCP-YA, Human Connectome Project Young Adult; SLEM, Southwest University Longitudinal Imaging Multimodal.

### Alignment of FC allometry with cortical hierarchy

We next asked whether regional variation in FC allometry follows the macroscale gradient of the human cortex. We examined the spatial correspondence between the FC allometry map and the canonical sensorimotor-to-association (S–A) axis, a principal gradient capturing unimodal-to-transmodal cortical hierarchy. The allometry map consistently showed significant negative correlations with the S–A axis across datasets (HCP-YA: *rho* = -0.626, *p*_spin_ < 0.001; SLEM: *rho* = -0.597, *p*_spin_ < 0.001; Figure 2), indicating that FC allometry is systematically organized along cortical hierarchy.

**Figure 2.**
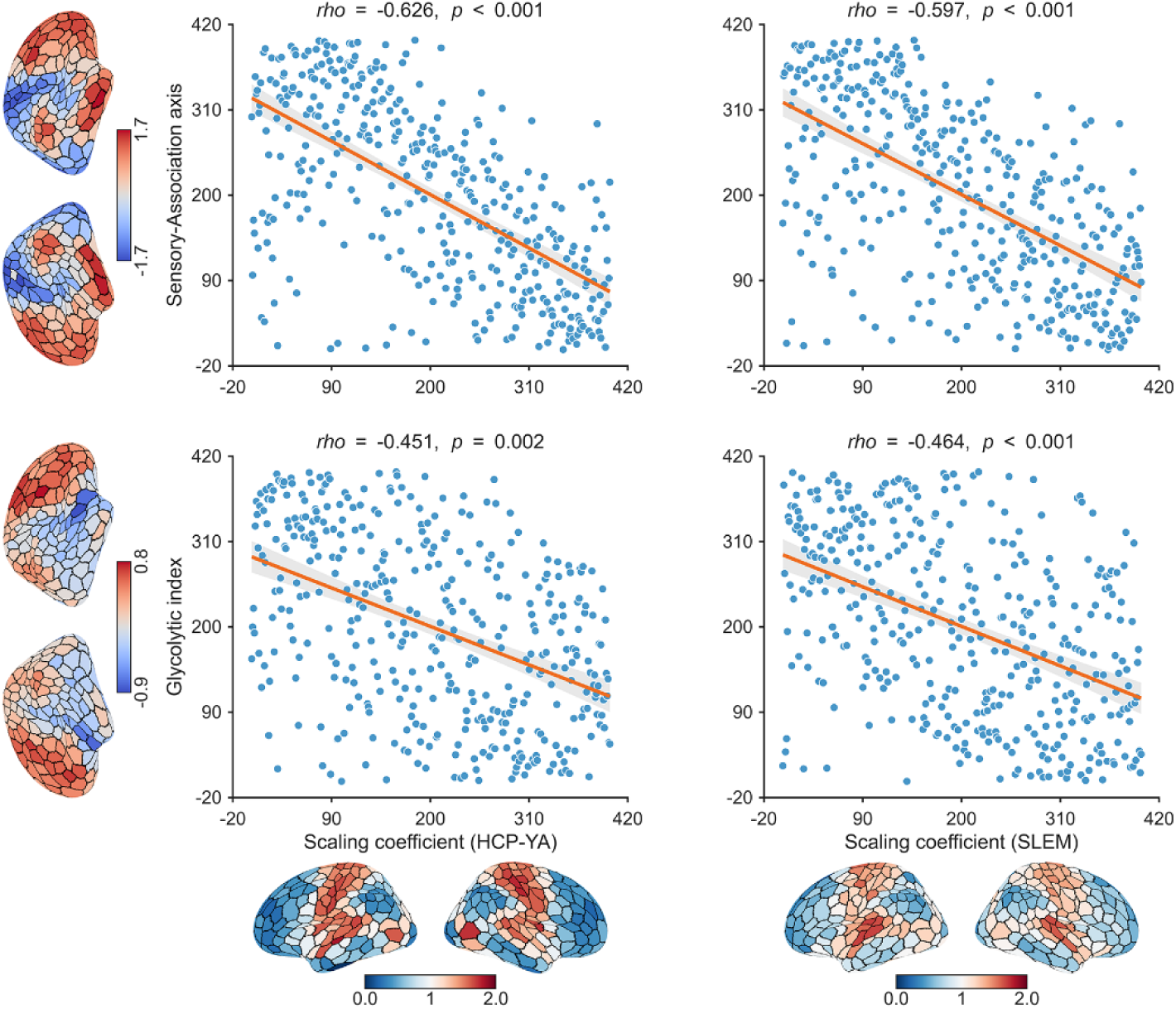
Relationships of FC allometry with cortical hierarchy and metabolic organization. Significant negative correlations were consistently observed between FC allometry and the S-A axis and glycolytic index map in both the HCP-YA and SLEM cohorts. HCP-YA, Human Connectome Project Young Adult; SLEM, Southwest University Longitudinal Imaging Multimodal.

Building on this global alignment, we further tested whether regions showing significant FC allometry were preferentially enriched within specific cortical modules. Cortical regions were assigned to predefined modules based on three complementary organizational schemes: (i) seven functional subnetworks defined by intrinsic FC [17], (ii) seven cytoarchitectonic classes derived from histological features [18], and (iii) four laminar classes reflecting levels of laminar differentiation [19]. Enrichment analysis revealed that regions of connectivity allometry followed a strongly non-random topographic distribution (*p*_spin_ < 0.05, FDR corrected).

The same modular enrichment pattern was observed across datasets (Figure 3). Specifically, regions showing positive allometry were significantly enriched in low-order systems, including the somatomotor subnetwork (HCP-YA: 76/114 regions, 66.7%; SLEM: 58/88 regions, 65.9%; both *p*_spin_ < 0.001), the primary motor cortex cytoarchitectonic class (HCP-YA: 23/114 regions, 20.2%; SLEM: 22/88 regions, 25.0%; both *p*_spin_ = 0.003), and the idiotypic laminar class (53/114 regions, 46.5%; SLEM: 43/88 regions, 48.9%; both *p*_spin_ < 0.001). By contrast, regions showing negative allometry were significantly enriched in high-order systems, including the default mode subnetwork (DMN; HCP-YA: 80/204 regions, 39.2%; SLEM: 77/159 regions, 48.4%; both *p*_spin_ < 0.001), the frontoparietal subnetwork (HCP-YA: 52/204 regions, 25.5%; SLEM: 43/159 regions, 27.0%; both *p*_spin_ < 0.001), and the heteromodal laminar class (HCP-YA: 110/204 regions, 53.9%; SLEM: 103/159 regions, 64.8%; both *p*_spin_ < 0.001). These findings indicate that FC allometry is tightly constrained by hierarchical and histological cortical architecture, with primary systems showing disproportionately greater connectivity expansion than transmodal systems.

**Figure 3.**
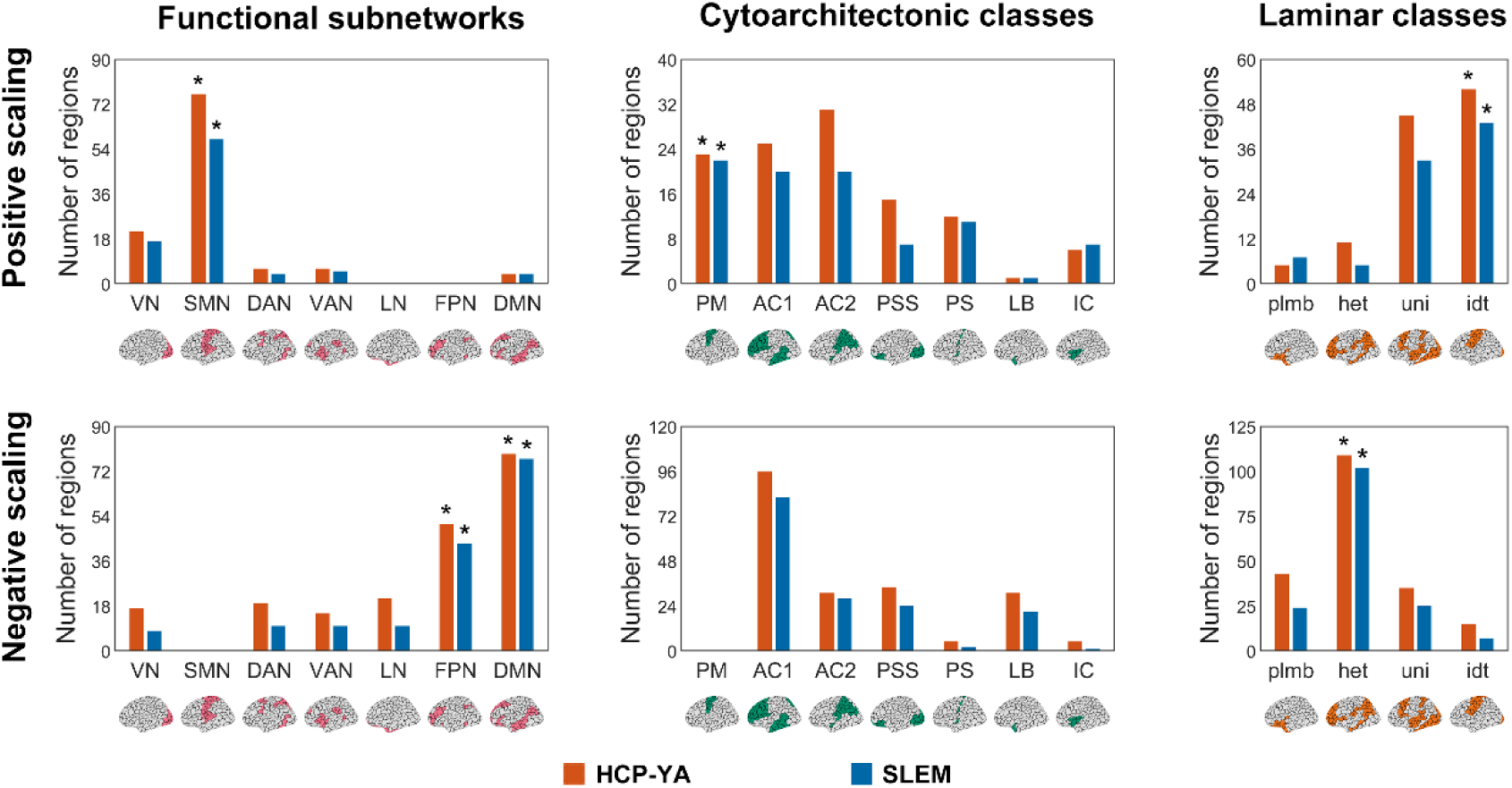
Distribution of FC allometry across cortical hierarchies. Across datasets, regions exhibiting significant positive allometry were consistently enriched in the somatomotor subnetwork, primary motor cortex cytoarchitectonic class, and idiotypic laminar class. By contrast, regions exhibiting significant negative allometry were consistently enriched in the frontoparietal and default mode subnetworks and heteromodal laminar class. VN, visual subnetwork; SMN, somatomotor subnetwork; DAN, dorsal attention subnetwork; VAN, ventral attention subnetwork; FPN, frontoparietal subnetwork; DMN, default mode subnetwork; PM, primary motor cortex; AC, association cortex; PSS, primary/secondary sensory cortex; PS, primary sensory cortex; LB, limbic regions; IC, insular cortex; plmb, paralimbic cortex; het, heteromodal cortex; uni, unimodal cortex; idt, idiotypic cortex; HCP-YA, Human Connectome Project Young Adult; SLEM, Southwest University Longitudinal Imaging Multimodal; *, *p*_spin_ < 0.05, FDR corrected.

### Metabolic substrate of FC allometry

Because FC is tightly coupled to metabolic demands [20–22], we next examined whether regional variation in FC allometry is related to the brain’s metabolic architecture. To do so, we tested the spatial correspondence between the FC allometry map and five publicly available metabolism-related maps, including cerebral blood flow, cerebral blood volume, oxygen metabolism, glucose metabolism, and glycolytic index [23]. Spatial autocorrelation was controlled via spin tests, and multiple comparisons were corrected via the FDR procedure.

Among these metabolic indices, the FC allometry map was consistently negatively correlated with the glycolytic index map across datasets (HCP-YA: *rho* = - 0.451, *p*_spin_ = 0.002; SLEM: *rho* = -0.464, *p*_spin_ < 0.001; Figure 2 and Table S1). This finding suggests that regions with higher aerobic glycolytic metabolism tend to exhibit lower scaling coefficients.

### Microscopic neurobiological signatures of FC allometry

To probe the molecular and cellular underpinnings of FC allometry, we examined its spatial correspondence with multiscale neurobiological maps, including 18 neurotransmitter receptor/transporter distributions [24] and 21 transcriptomic profiles indexing specific cell types, biological processes, and developmental stages [25]. Spatial autocorrelation was controlled via spin tests, and multiple comparisons were corrected via the FDR procedure.

Among the neurotransmitter maps, the FC allometry map consistently showed significant positive correlations with the noradrenaline transporter (NET) and vesicular acetylcholine transporter (VAChT) maps and significant negative correlations with the serotonin receptor 2a (5-HT_2a_) map (Figure 4 and Table S2). For the transcriptomic profiles, the FC allometry maps consistently showed significant positive correlations with average expression of genes enriched in oligodendrocytes, axon development, and myelination, and significantly negative correlations with the average expression of genes enriched in astrocytes and cortical developmental stages of neonatal early infancy and early childhood (Figure 4 and Table S3). In addition, the FC allometry map derived from the SLEM dataset showed several dataset-specific correlations, including negative correlations with serotonin receptor 4 (5-HT_4_) and μ-opioid receptor (MOR) maps, as well as with genes related to excitatory neurons, inhibitory neurons, synapse development, and cortical developmental stages of middle-to-late childhood (Figure 4 and Table S3).

**Figure 4.**
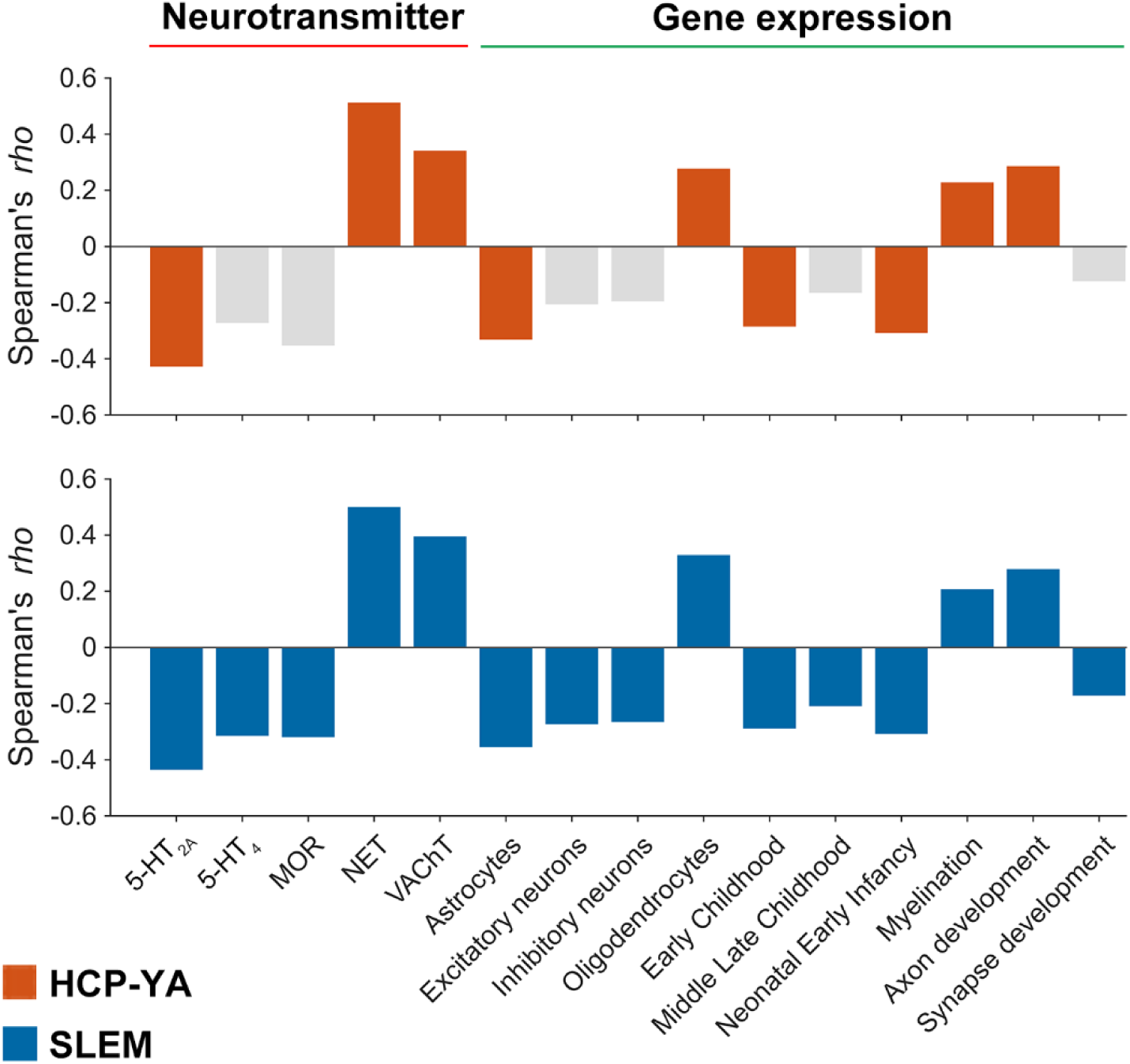
Neurobiological correlates of FC allometry. Bar plots show spatial correlations between FC allometry and neurotransmitter receptor/transporter distributions and transcriptomic profiles related to specific cell types, developmental stages, and biological processes in the HCP-YA and SLEM cohorts. Colored bars indicate significant associations, whereas gray bars indicate non-significant associations. HCP-YA, Human Connectome Project Young Adult; SLEM, Southwest University Longitudinal Imaging Multimodal; 5HT_2a_, serotonin receptor 2a; 5-HT_4_, serotonin receptor 4; MOR, μ-opioid receptor; NET, noradrenaline transporter; VAChT, vesicular acetylcholine transporter.

### Functional significance of FC allometry

We further asked whether regional variation in FC allometry relates to behavior and cognition. To answer this question, we correlated the FC allometry map with 123 meta-analytic activation maps from the NeuroSynth database. The behavioral and cognitive terms related to these activation maps are listed in Table S4. Spatial autocorrelation was controlled via spin tests, and multiple comparisons were corrected via the FDR procedure.

The FC allometry map derived from the HCP-YA dataset exhibited significant positive and negative correlations with 18 and 42 meta-analytic activation maps, respectively (*p*_spin_ < 0.05, FDR corrected). For the SLEM dataset, the FC allometry map showed significant positive and negative correlations with 19 and 41 meta-analytic activation maps, respectively (*p*_spin_ < 0.05, FDR corrected). These functional associations were highly similar across the two datasets (Dice coefficient = 0.967). Positive correlations were mainly related to lower-order functions, such as sensory (e.g., ‘multisensory’), perception (e.g., ‘perception’ and ‘visual perception’), and motor (e.g., ‘motor control’, ‘movement’, ‘coordination’, and ‘rhythm’). By contrast, negative correlations were primarily associated with higher-order functions, including executive function (e.g., ‘cognitive control’ and ‘strategy’), memory (e.g., ‘memory retrieval’, ‘episodic memory’, and ‘autobiographical memory’), and decision-making (e.g., ‘reasoning’, ‘judgment’, ‘decision’, and ‘decision making’) (Figure 5 and Table S5).

**Figure 5.**
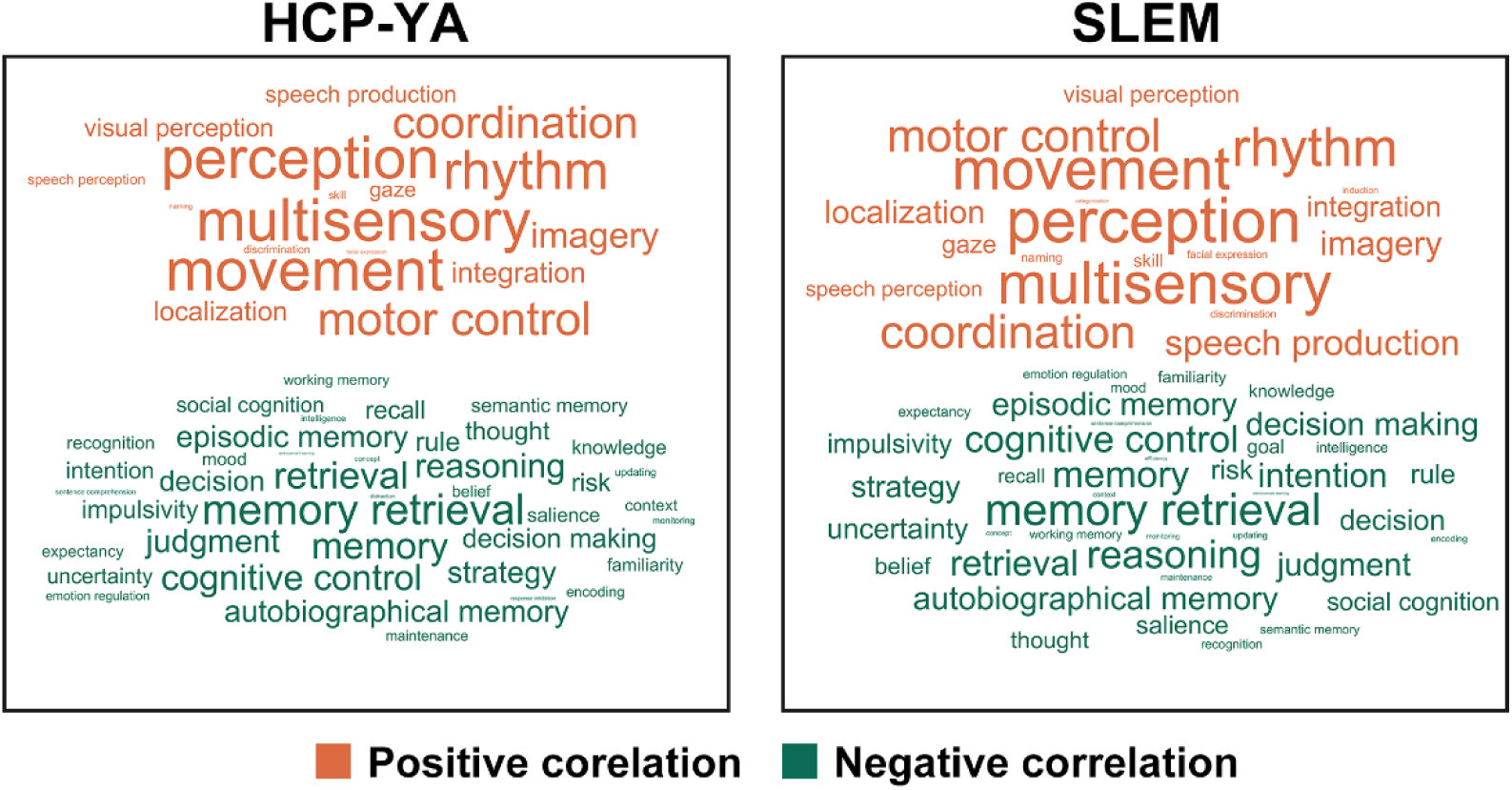
Cognitive associations of FC allometry. Word clouds illustrate cognitive terms significantly associated with FC allometry in the HCP-YA and SLEM cohorts. Word size represents the absolute correlation magnitude between FC allometry and Neurosynth meta-analytic activation maps. HCP-YA, Human Connectome Project Young Adult; SLEM, Southwest University Longitudinal Imaging Multimodal.

### State-dependent reconfiguration of FC allometry

Although our functional decoding analysis suggested links between FC allometry and cognitive function, it remains unclear whether this scaling principle is modulated by active task engagement. To address this question, we first derived a FC allometry map from the HCP-YA task dataset, in which participants performed a working memory task. The task-derived allometry map was highly similar to the map derived from the HCP-YA resting-state dataset (*rho* = 0.661, *p*_spin_ < 0.001; Figure 6A), indicating that the core principle of FC allometry is largely state-invariant.

**Figure 6.**
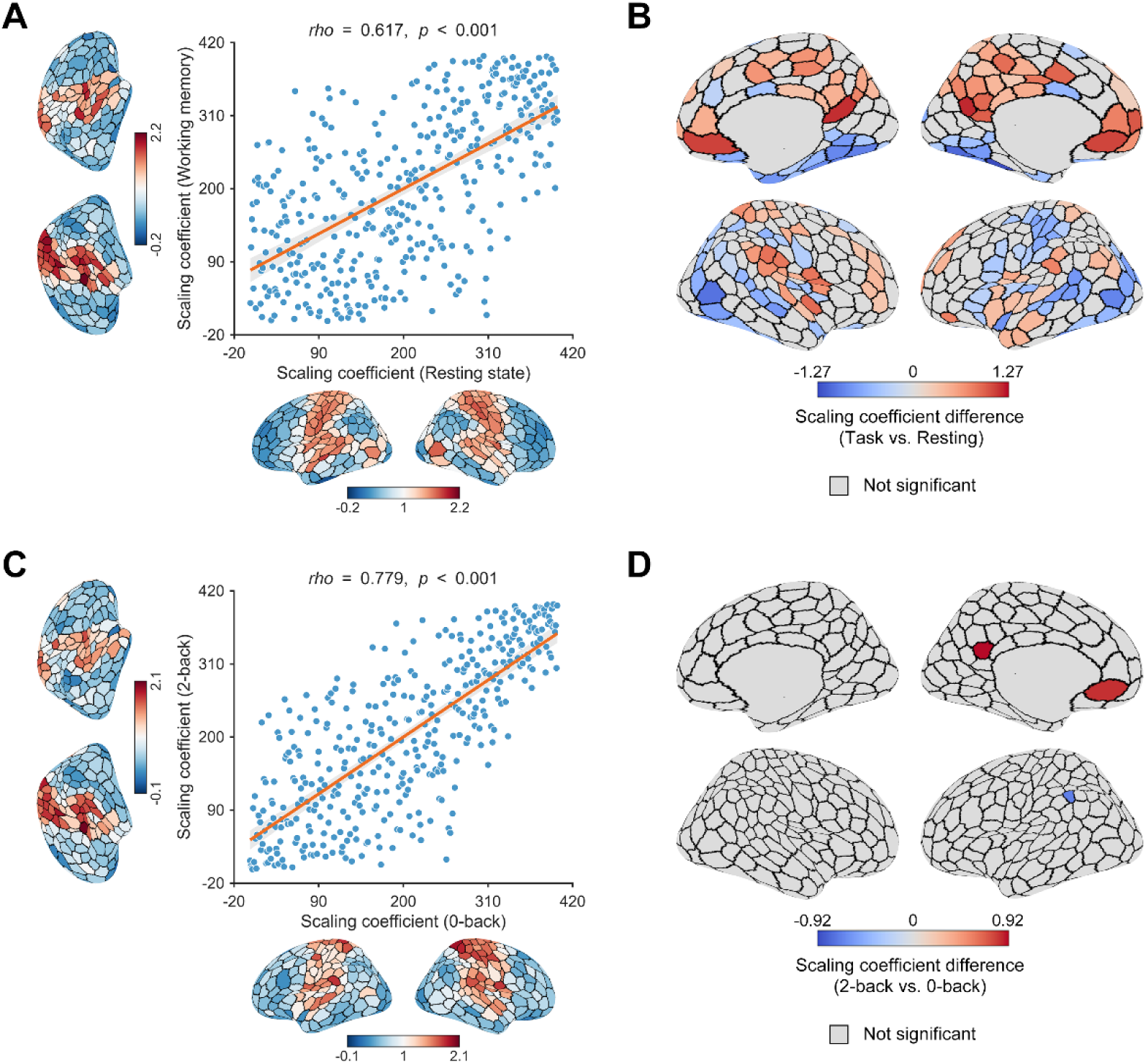
State-dependent modulation of FC allometry during working memory performance. **A**) A high positive correlation was observed between FC allometry maps derived from resting-state and working memory task conditions. **B**) A total of 194 regions were identified to show significant differences in the scaling coefficients between the task and resting-state conditions. **C**) A high positive correlation was observed between FC allometry maps derived from 0-back and 2-back conditions. **D**) Compared with the 0-back condition, the 2-back condition was associated with significant alterations in the scaling coefficients of three regions.

Despite this global stability, localized reconfiguration was evident during task performance. A within-subject permutation test (10,000 iterations) identified 194 regions showing significant differences between rest and task (all *p* < 0.05, FDR corrected), including 98 regions with increased scaling coefficients and 96 with decreased coefficients during task engagement (Figure 6B). The increased regions were primarily located in the frontal pole, the precuneus, and the cingulate gyrus, whereas the decreased regions were predominantly located in the occipital lobe, the ventral temporo-occipital cortex, and the fusiform gyrus. Enrichment analysis revealed that the regions showing decreased scaling coefficients were preferentially located in the ventral attention subnetwork (VAN; 23/98 regions, 23.5%, *p* = 0.002).

We further investigated whether FC allometry is sensitive to cognitive demands by comparing 0-back and 2-back conditions. The corresponding allometry maps were highly similar between conditions (*rho* = 0.812, *p*_spin_ < 0.001; Figure 6C), indicating that the allometric principle remains robust across varying levels of cognitive load. Nevertheless, increasing task load was associated with a decreased scaling coefficient in the left supramarginal gyrus within the frontoparietal subnetwork, whereas scaling coefficients increased in two DMN nodes located in the left ventromedial prefrontal and paracingulate cortices (*p* < 0.05, FDR corrected; Figure 6D).

Taken together, these findings indicate that FC allometry is a stable organizational property across mental states, while retaining the flexibility for fine-grained, region-specific modulation in response to cognitive demands.

### Global preservation but regional dysregulation of FC allometry in ASD

Given our finding that FC allometry was linked to gene expression profiles associated with early developmental stages, we next examined whether this organizing principle is altered in ASD, a neurodevelopmental condition characterized by atypical FC. To establish an appropriate developmental baseline, we first tested whether FC allometry was already present in the juvenile brain using the ABIDE dataset (ages 8–18). The allometry map derived from typically developing controls (TDCs) in the ABIDE dataset showed strong spatial correspondence with the FC allometry map derived from adult cohorts (HCP-YA: *rho* = 0.661, *p*_spin_ < 0.001; SLEM: *rho* = 0.734, *p*_spin_ < 0.001), indicating that this scaling architecture is already established by late childhood.

Against this developmental baseline, we then examined FC allometry in ASD. At the global level, the ASD allometry map remained highly similar to that of TDCs (*rho* = 0.558, *p*_spin_ < 0.001; Figure 7A), suggesting that the overall pattern of FC allometry is largely preserved in the disorder. However, regional analysis using a permutation framework (10,000 iterations) identified 10 regions that showed significant alterations in the scaling coefficients in ASD (all *p* < 0.05, FDR corrected; Figure 7B). Of these, eight regions showed increased scaling coefficients in ASD, primarily localized in core nodes of the DMN, including the precuneus, posterior cingulate cortex, and medial prefrontal cortex. By contrast, decreased scaling coefficients were confined to two regions within the somatomotor subnetwork, specifically the pericentral and opercular areas. These findings indicate that, despite preservation of the global allometric framework, ASD is associated with region-specific dysregulation of FC allometry, characterized by disproportionately greater scaling in higher-order association cortex and reduced scaling in primary sensorimotor regions.

**Figure 7.**
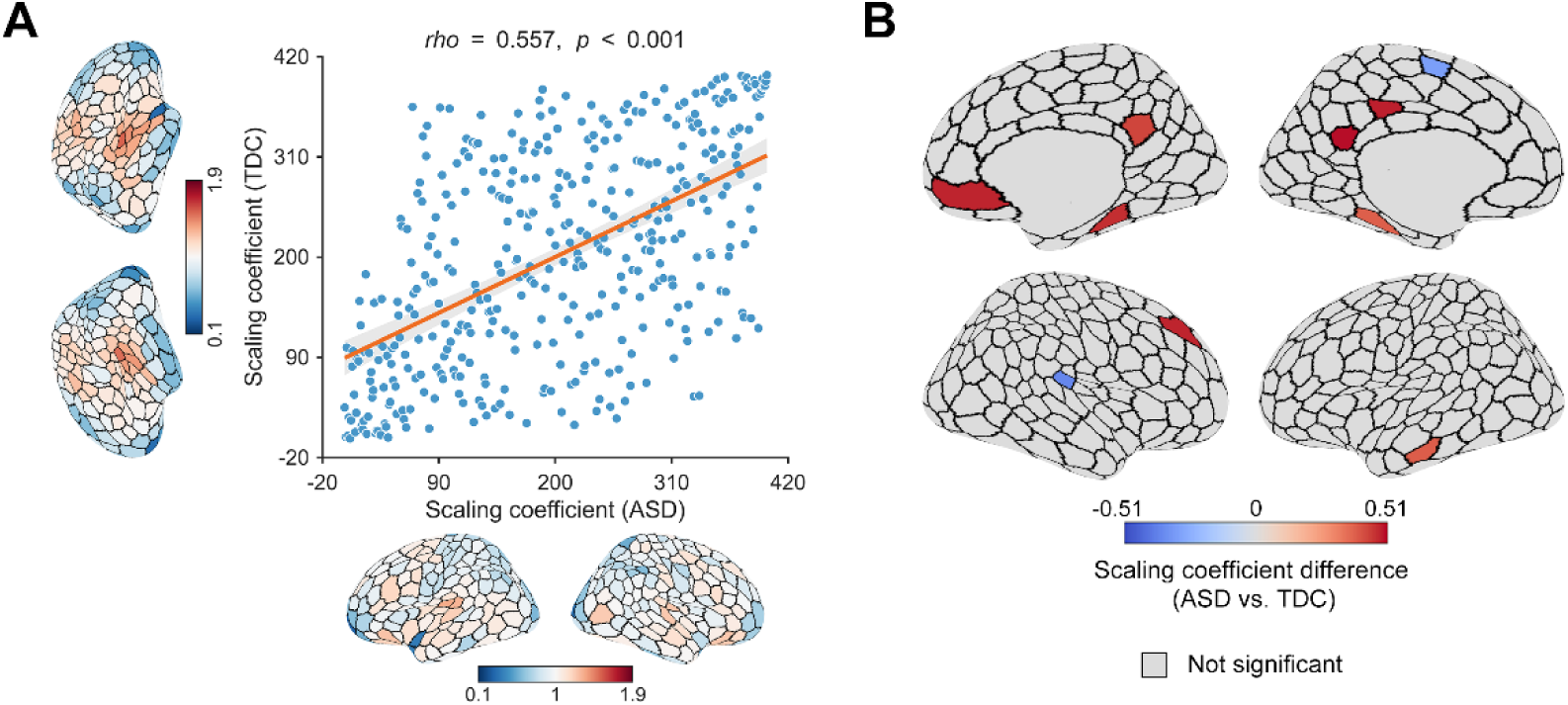
Altered FC allometry in ASD. **A**) A high positive correlation was observed in the scaling coefficients between the ASD and TDC groups. **B**) Compared with the TDC group, the ASD group showed significantly increased scaling coefficients in eight regions and decreased scaling coefficients in two regions. ASD, autism spectrum disorder; TDC, typically developing control.

## Materials and Methods

### Participants and MRI data acquisition

This study included four datasets. For each dataset, the participant recruitment processes and informed consent forms were ratified by the corresponding Institutional Review Board. This study was approved by the Institutional Review Board of the Institute for Brain Research and Rehabilitation at South China Normal University.

#### HCP-YA resting-state dataset

Our discovery cohort consisted of 413 unrelated participants from the HCP-YA dataset. All scans were acquired using a customized 3T scanner at Washington University in St. Louis. Structural MRI were acquired using a magnetization-prepared rapid gradient-echo sequence with the following parameters: repetition time (TR) = 2,400 ms, echo time (TE) = 2.14 ms, flip angle (FA) = 8°, 256 slices, matrix = 320 × 320, field of view (FOV) = 224 × 224 mm^2^, slice thickness/gap = 0.7/0 mm, and voxel size = 0.7 × 0.7 × 0.7 mm^3^. Resting-state fMRI data were obtained using a multiband echo-planar imaging (EPI) sequence with the following parameters: TR = 720 ms, TE = 33 ms, FA = 52°, 72 slices, matrix = 104 × 90, FOV = 208 × 180 mm^2^, slice thickness/gap = 2/0 mm, and voxel size = 2 × 2 × 2 mm^3^. Each participant completed two resting-state sessions on consecutive days, and each session included two runs acquired with left-to-right and right-to-left phase-encoding directions. Only the left-to-right run from the first session was included in the present analyses.

#### SLEM dataset

The SLEM dataset included 580 healthy participants at baseline, of whom 240 returned for a second visit and 228 for a third visit. For the present study, we included 464 participants who completed both structural MRI and resting-state fMRI scans at baseline. All MRI scans were acquired on a Siemens Trio 3 T scanner at Southwest University Center for Brain Imaging. Structural MRI scans were acquired using a magnetization-prepared rapid gradient-echo sequence with the following parameters: TR = 1,900 ms, TE = 2.52 ms, FA = 9°, matrix = 256 × 256, FOV = 256× 256 mm^2^, 176 slices, slice thickness/gap = 1/0 mm, and voxel size = 1 × 1 × 1 mm^3^. Resting-state fMRI scans were obtained using a gradient EPI sequence with the following parameters: TR = 2,000 ms, TE = 30 ms, FA = 90°, FOV = 220 × 220 mm^2^, 32 slices, slice thickness/gap = 3/1 mm, and voxel size = 3.4 × 3.4 × 3 mm^3^.

#### HCP-YA task dataset

To assess task-related modulation of FC allometry, we analyzed a subset of 400 participants from the HCP-YA discovery cohort who completed a working-memory task. The acquisition parameters for task-based fMRI were identical to those used for the resting-state scans. The task followed an N-back paradigm. Each run comprised eight 25-s blocks, equally divided between 0-back and 2-back conditions. Stimuli consisted of four categories (i.e., faces, tools, places, and body parts) with each category presented in two blocks per run. Each block was preceded by a 2.5-s cue indicating the task condition and stimulus category [26].

#### ABIDE dataset

To evaluate the clinical relevance of FC allometry, we utilized a large-scale multisite dataset from the ABIDE I and II cohorts [27,28]. The final sample included 338 individuals with ASD and 518 TDCs aggregated from 20 sites. Participants were included according to the following criteria: (i) age between 8 and 18 years; (ii) controlled head motion, defined as mean framewise displacement ≤ 0.5 mm or maximum framewise displacement ≤ 3 mm; and (iii) absence of substantial visual artifacts. Sites contributing fewer than five participants per group were further excluded. To mitigate potential multisite effects, we applied ComBat harmonization to the resulting FC data [29], while preserving biological variability related to age, sex, and diagnostic group. The final ASD and TDC groups were well matched for age, sex, head motion, full-scale IQ, handedness, and acquisition site (Table S6). Detailed diagnostic procedures, ethical statements, and scanning parameters are publicly accessible through the ABIDE website (https://fcon_1000.projects.nitrc.org/indi/abide/).

### FMRI data preprocessing

FMRI data preprocessing was implemented using standardized pipelines matched to each data source while maintaining methodological consistency wherever possible. For the HCP-YA resting-state and task datasets, we used data that had already undergone the HCP minimal preprocessing pipeline, including gradient distortion correction, motion correction, field map-based EPI distortion correction, registration of functional images to structural images, nonlinear registration to the Montreal Neurological Institute (MNI) space, and grand-mean intensity normalization [30,31].

The preprocessed fMRI data were then band-pass filtered (0.01–0.08 Hz), and nuisance covariates were further regressed out, including 24 parameter head motion profiles, white matter signals, cerebrospinal fluid signals, and global signals. Filtering and nuisance regression were implemented in a single regression model to avoid reintroducing artifacts [32].

For the SLEM and ABIDE datasets, functional images were preprocessed using the GRETNA toolbox [33] based on the SPM12 package (http://www.fil.ion.ucl.ac.uk/spm/software/spm12). This involved removal of the first five volumes, slice-timing correction using Sinc interpolation, and head-motion correction using rigid-body transformation. Afterwards, band-pass filtering and nuisance regression were performed in the same manner as for the HCP-YA datasets. Finally, functional images were spatially normalized to the MNI space by applying the transformation fields derived from tissue segmentation of the corresponding structural images.

### Network construction

In this study, we utilized a functionally defined atlas to parcellate the cerebral cortex into 400 regions [17]. For each participant in the HCP-YA resting-state, SLEM, and ABIDE datasets, the mean time series of each region was extracted by averaging the fMRI signals across all voxels within that region. Inter-regional FC was then estimated by computing the Pearson correlation coefficient between each pair of regional mean time series. Negative correlations were excluded from subsequent analyses.

For the HCP-YA task dataset, we first extracted the mean time series of each region from the full task session. We then partitioned the task time series into 0-back and 2-back segments based on the experimental block design. To account for the haemodynamic response delay, we applied a 6-s shift (8 TRs) and concatenated blocks from the same condition to reconstruct condition-specific time series. Inter-regional FC was subsequently calculated for the full task session as well as for the 0-back and 2-back time series using Pearson correlation. This procedure yielded three functional brain networks per participant in the HCP-YA task dataset. Negative correlations were again excluded from further analyses.

### Nodal and global FC strength

For each functional brain network constructed above, we defined *S*_global_ as the total FC strength across all edges in the network, and *S*_nodal_ for each region as the total FC strength across all edges linking that region to the other regions in the network.

### Estimation of FC allometry

To estimate the scaling relationship between *S*_nodal_ and *S*_global_, we fitted the following log–log regression model for each brain region:

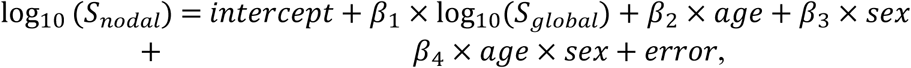

where *β*_1_ denotes the scaling coefficient. A value of *β*_1_ = 1 indicates isometric scaling, such that *S*_nodal_ changes proportionally with *S*_global_. By contrast, deviation from unity indicates allometric scaling: *β*_1_ > 1 indicates that *S*_nodal_ increases faster than *S*_global_ (i.e., positive allometry), whereas *β*_1_ < 1 indicates that *S*_nodal_ increases more slowly than *S*_global_ (i.e., negative allometry).

To test whether the observed scaling coefficient differed significantly from isometry, we calculated a *t*-statistic as:

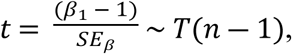

where *SE_β_* is the standard error of *β*_1_ estimated via bootstrapping. Statistical significance was evaluated against a *t*-distribution with n-1 degrees of freedom, where n is the number of participants. Multiple comparisons across all brain regions were corrected using the FDR procedure at *q* < 0.05.

Because exact ages were unavailable in the HCP-YA dataset, age was treated as a categorical variable in the corresponding regression models. To examine potential sex differences in FC allometry, we additionally fitted the log–log regression model separately in male and female participants for the HCP-YA resting-state and SLEM datasets:

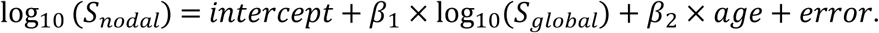

### Relationship between FC allometry and cortical topography

To characterize the spatial organization of FC allometry, we performed two complementary analyses to examine whether regional variation in scaling coefficients was systematically related to cortical hierarchy: global alignment and modular enrichment.

#### Global alignment

We first tested whether the FC allometry followed the macroscale gradient of human cortical organization. Specifically, we quantified the spatial correspondence between regional scaling coefficients and the S-A axis derived from Neuromaps [24], a principal gradient capturing the primary-to-transmodal hierarchical organization of the cortex [34], using Spearman rank correlation. Statistical significance was assessed with a spin-test framework, which controls spatial autocorrelation by randomly rotating brain maps on cortical surface [35]. We generated 10,000 random spatial rotations (i.e., spins) of the FC allometry map and, for each rotation, recalculated its correlation with the S-A axis. A two-sided *P*-value was calculated as the proportion of spins for which the absolute correlation coefficient was greater than or equal to the observed absolute value. This procedure preserved the spatial covariance structure of the cortical map and thus accounted for potential confounding by spatial autocorrelation.

#### Modular enrichment

Complementing the global alignment, we next tested whether regions showing significant positive or negative FC allometry were preferentially enriched within specific cortical modules. Cortical regions were assigned to canonical modules according to three complementary organizational schemes: 1) functional subnetworks, consisting of seven modules defined by intrinsic FC (visual, somatomotor, dorsal attention, VAN, limbic, frontoparietal, and DMN) [36]; 2) cytoarchitectonic classes, comprising seven modules based on histological features (primary motor cortex, association cortex 1, association cortex 2, primary/secondary sensory, primary sensory cortex, limbic regions, and insular cortex) [18]; and 3) laminar classes, comprising four modules with different levels of laminar differentiation (idiotypic, unimodal, heteromodal, and paralimbic) [19]. Modular assignments for the 400 cortical regions were taken from previous studies [17,37,38]. For each module, we counted the number of regions showing significant positive or negative allometry within it. Statistical significance was again assessed using a spin-test framework by rotating the FC allometry map (10,000 spins) to construct null distributions for the observed numbers expected in each module by chance. Multiple comparisons were controlled using the FDR procedure within each organizational scheme at *q* < 0.05.

### Relationship between FC allometry and cortical metabolism

Previous studies have found that highly connected regions are associated with high rates of energy metabolism [20,21,23,39]. Thus, we examined whether regional variation in FC allometry were related to regional differences in energy metabolism. Specifically, we calculated Spearman rank correlations between the FC allometry map and five publicly available metabolism-related maps, including cerebral blood flow, cerebral blood volume, oxygen metabolism, glucose metabolism, and glycolytic index [23]. Statistical significance was assessed using spin tests to account for spatial autocorrelation (10,000 spins).

### Relationship between FC allometry and microscopic cortical organization

To probe the molecular signatures underlying regional variation in FC allometry, we calculated Spearman rank correlations between the FC allometry map and two classes of cortical maps: neurotransmitter maps and gene-expression maps. Statistical significance was assessed using spin tests to account for spatial autocorrelation (10,000 spins), and multiple comparisons were controlled using the FDR procedure at *q* < 0.05.

#### Neurotransmitter maps

Neurotransmitter maps were obtained from the Neuromaps toolbox [24], which includes 25 positron emission tomography maps covering 18 receptors and transporters. These comprised the 5-HT_2a_, 5-HT_4_, serotonin receptors 1a, 1b, and 6, serotonin transporter, dopamine receptors 1 and 2, dopamine transporter, α_4_β_2_ nicotinic acetylcholine receptor, muscarinic acetylcholine receptor M1, VAChT, metabotropic glutamate receptor 5, cannabinoid receptor 1, γ-aminobutyric acid receptor A, MOR, NET, and histamine receptor 3. For targets with multiple available tracers, standardized maps were generated by calculating the weighted average across studies. Region-wise mean intensities were extracted for spatial correlation analyses.

#### Gene-expression maps

Gene-expression data were obtained from the Allen Human Brain Atlas dataset [25], a publicly available resource comprising brain-wide transcriptional data from six healthy adult donors (five males; 24–57 years) with no known history of neurological or psychiatric disease. Transcriptional activity was measured for 20,737 genes across 3,702 spatially distinct tissue samples spanning nearly the entire brain. These data were processed using the abagen toolbox [40] according to established recommendations [41]. From these data, we screened gene sets enriched for seven canonical cell types (excitatory neurons, inhibitory neurons, oligodendrocyte progenitor cells, astrocytes, endothelial cells, microglia and oligodendrocytes) [41], four neurodevelopmental processes (axon development, myelination, dendrite development, and synapse development) [42], and ten developmental stages from early fetal to young adult [43]. For each category, mean regional expression levels were computed and then averaged across donors, yielding a total of 21 gene-expression maps.

### Functional decoding of FC allometry

To assess the cognitive relevance of regional variation in FC allometry, we examined the spatial correspondence between the FC allometry map and with a set of meta-analytic activation maps. The activation maps were obtained from NeuroSynth (https://github.com/neurosynth/neurosynth), which quantifies associations between brain voxels and terms of neurocognitive processes [44]. Specifically, NeuroSynth estimates, for each voxel, the probability that a given term is reported in a neuroimaging experiment if neural activation is observed at that voxel. Because our focus was on cognition and behavior, we restricted the analysis to 123 cognitive and behavioral terms based on the Cognitive Atlas [45], a public ontology of cognitive science that includes a comprehensive list of neurocognitive processes. The resulting term-specific association maps were parcellated into 400 regions and z-scored. For each association map, we calculated its Spearman rank correlation with the FC allometry map. Statistical significance was estimated using spin tests to account for spatial autocorrelation (10,000 spins). Multiple comparisons across all association maps were controlled using the FDR procedure at *q* < 0.05.

### Task-related modulation of FC allometry

To determine whether FC allometry is preserved during active cognitive engagement, we first assessed the spatial consistency between the allometry map derived from the full task session in the HCP-YA task dataset and that derived from the HCP-YA resting-state dataset using Spearman rank correlation. Statistical significance was estimated using spin tests to account for spatial autocorrelation (10,000 spins). We then compared regional scaling coefficients between the resting-state and full-task allometry maps using a within-subject permutation framework (10,000 iterations), with state labels randomly shuffled within each participant to generate a null distribution. To further examine the effects of cognitive load, we compared regional scaling coefficients between the 2-back and 0-back conditions using the same within-subject permutation procedure, with condition labels randomly shuffled within each participant (10,000 iterations).

### Alterations of FC allometry in ASD

To determine whether FC allometry is preserved or disrupted in ASD, we first assessed the spatial consistency between the FC allometry maps derived from the ASD and TDC groups using Spearman rank correlation. Statistical significance was estimated using spin tests to account for spatial autocorrelation (10,000 spins). To identify localized alterations, we then used a permutation-based approach to test for between-group differences in regional scaling coefficients by randomly shuffling diagnosis labels across participants (10,000 iterations).

## Discussion

In this study, we identified a fundamental allometric scaling law of the human functional connectome. Rather than responding uniformly to global FC fluctuations, cortical regions exhibited substantial variability in the extent to which their FC scaled with global dynamics. Regional scaling coefficients were systematically organized along the S–A axis, with primary sensory and transmodal association regions anchoring opposite ends of the allometric spectrum. This architecture was robust across sexes and age groups, closely linked to metabolic and molecular substrates, and replicated across independent cohorts from different cultural backgrounds. Furthermore, although localized reconfigurations emerged during cognitive task states and in ASD, the overall allometric pattern remained largely preserved. Collectively, these findings establish FC allometry as a fundamental organizing principle of the human brain, one that is aligned with cortical hierarchy, rooted in biological substrates, and stable across populations and brain states.

### Robust hierarchical organization of FC allometry

A central finding of this study was that FC allometry was not randomly distributed across the cortex, but instead followed a highly organized spatial pattern aligned with the S–A axis. Specifically, the global topography of FC allometry showed a strong correspondence with this axis, with significant positive allometry concentrated in lower-order systems and negative allometry concentrated in higher-order systems. This interpretation was further supported by meta-analytic decoding, which showed that the FC allometry map correlated positively with sensory–motor activation maps and negatively with maps related to executive and mnemonic functions. We suggest that this divergent topography may reflect an alignment between functional demands and the biological constraints under which different cortical systems operate. Low-order regions are primarily responsible for processing sensory input and executing motor output. To support these specialized functions, they rely predominantly on localized, short-range connections [46,47]. The relatively low metabolic cost of such wiring may provide these regions with greater economical flexibility, allowing their FC to fluctuate more strongly at lower physiological cost, thereby manifesting as positive allometry. By contrast, high-order regions act as integrative centers and network hubs of the brain [48]. Supporting large-scale information integration requires stable, long-range, and metabolically costly connectivity profiles spanning distributed brain systems [46,47,49]. These requirements may constrain the extent to which high-order regions can vary proportionally with global fluctuations. As a result, high-order regions remain relatively less coupled to global variation, giving rise to the observed negative allometry. From this perspective, the spatial pattern of FC allometry may reflect a macroscale trade-off between minimizing wiring cost and maximizing communication efficiency in the brain [50,51]. This interpretation was further supported by the observed correlation between FC allometry and the regional glycolytic index.

Importantly, this hierarchical organization does not appear to be transient or cohort-specific, but instead represents a stable intrinsic feature of brain organization. We found that the spatial distribution of FC allometry was highly consistent across sexes, age groups, cultures, task states, and clinical conditions. Such reproducibility strongly supports the view that FC allometry represents an intrinsic organizational architecture of the human brain that is largely independent of demographic variation and state-related effects.

### Biological substrates underlying FC allometry

We found that FC allometry was closely related to the spatial distribution of specific genes and neurotransmitters, supporting the view that macroscale functional architecture is strongly constrained by molecular and chemoarchitectonic landscape [52,53]. At the transcriptomic level, FC allometry showed significant correlations with genes enriched during infancy and childhood. This developmental signature is consistent with the observation that the spatial pattern of FC allometry mirrors the known temporal sequence of cortical maturation, whereby primary sensory–motor systems exhibit adult-like topology at birth, whereas higher-order association cortices continue to mature throughout adolescence [54]. Together with our finding that adult-like FC allometry is already evident in children, these results suggest that the scaling architecture of FC may be established early in development and may serve as a scaffold for subsequent cortical maturation. In addition, FC allometry showed strong spatial correspondence with genes enriched in oligodendrocytes, myelination, and axon development. Collectively, these processes reflect the formation and maintenance of myelin sheaths by oligodendrocytes around developing axons [55], which provide both structural support and metabolic efficiency for rapid neural signal conduction [56]. Accordingly, our results suggest that white-matter infrastructure may provide a physiological scaffold for FC allometry. More specifically, its signaling capacity and metabolic support may define the physical constraints within which regional FC can scale.

Beyond transcriptomic associations, the FC allometry map also showed significant spatial correspondence with specific neurotransmitter systems, including the NET, VAChT and 5-HT_2a_ receptor. Among these, the association with the 5-HT_2a_ is particularly notable given the established role of this receptor in shaping macroscale cortical functional organization [57,58]. At the cellular level, 5-HT_2a_ receptor is highly expressed on layer 5 pyramidal neurons, which are key elements facilitating long-range corticocortical projections and global brain dynamics [59–63]. Functionally, 5-HT_2a_ signaling has been shown to modulate cortical hierarchy by promoting communication across networks that are typically segregated [64], potentially conferring greater flexibility to large-scale connectivity dynamics. The spatial alignment between 5-HT_2a_ distribution and FC allometry therefore suggests that serotonergic system may contribute to the neurochemical tuning underlying the brain’s macroscopic allometric architecture.

Taken together, these results suggest that FC allometry is jointly supported by at least two complementary biological systems: the structural scaffold provided by white matter and the dynamic regulation provided by neuromodulatory systems. In this framework, the former may constrain the feasible range of FC scaling, whereas the latter may fine-tune its expression across cortical systems.

### Cognitive and clinical implications of FC allometry

Previous studies have shown that FC is dynamically modulated by cognitive demands [65,66] and is disrupted in neurodevelopmental disorders [67–69]. Against this background, our findings are notable in showing that the allometric scaling rules governing FC remain remarkably stable across both mental states and clinical conditions. This stability suggests that FC allometry reflects an intrinsic and robust organizing principle of the human brain. Nevertheless, this global preservation does not preclude local flexibility. Rather, our results indicate that FC allometry combines a stable large-scale backbone with context- and disorder-sensitive regional reconfigurations.

During working-memory task performance, we observed widespread regional reconfiguration of FC allometry, characterized by increased scaling coefficients in executive regions and decreased coefficients in occipito-temporal and fusiform areas. Notably, enrichment analysis revealed that regions with decreased scaling coefficients were preferentially located within the VAN. Previous studies have shown that the VAN is suppressed during working-memory performance to prevent attentional reorienting toward task-irrelevant stimuli [70]. Our results extend this view by suggesting that task-related VAN suppression may involve not only reduced activation, but also altered scaling with respect to global FC fluctuations. Specifically, the decreased scaling coefficients imply that these regions become less sensitive to whole-brain FC changes during working-memory task performance, consistent with a form of functional decoupling from global dynamics. In this sense, FC allometry may provide a systems-level mechanism through which the brain shields task-relevant processing from interference by globally shared fluctuations.

A similar dissociation between global stability and local dysregulation was observed in clinical conditions. Although the overall FC allometric architecture was preserved in ASD, we identified 10 regions showing significant allometric abnormalities. Strikingly, the most prominent alterations involved increased scaling coefficients in core DMN nodes, including the precuneus, posterior cingulate cortex, and medial prefrontal cortex. Previous studies have reported increased local and within-network connectivity in the DMN in ASD [71]. Our findings suggest that such hyperconnectivity may be related to the increased FC allometry observed in this study. If global FC tends to increase during development, particularly across adolescence [72], then elevated scaling coefficients in ASD may render DMN regions disproportionately sensitive to global fluctuations. This increased sensitivity could amplify coordinated DMN coupling beyond the normative range, thereby contributing to the overconnectivity pattern repeatedly reported in ASD. More broadly, these findings raise the possibility that FC alterations in brain disorders may arise not only from altered absolute connection strength, but also from altered scaling rules that determine how local regions are embedded within global network dynamics.

Taken together, these results suggest that FC allometry may have both cognitive and clinical relevance. Cognitively, it appears to support adaptive reconfiguration by selectively modulating the degree to which regional FC tracks global dynamics during task engagement. Clinically, its local disruption, despite preserved global architecture, may provide a mechanistic link between macroscale network organization and disorder-related dysconnectivity. This combination of global stability and local plasticity may be a key feature of how the brain balances robustness with functional specialization across both healthy and pathological states.

### Limitations and future directions

Several limitations should be acknowledged. First, our analyses focused exclusively on the cerebral cortex. Since subcortical and cerebellar structures play essential roles in whole-brain coordination, future work should incorporate these regions to provide a more comprehensive account of FC allometry across the entire brain. Second, we identified multiple molecular signatures associated with regional variation in the FC allometry by leveraging several publicly available datasets. Because these datasets were derived from different populations, the molecular correlates reported here may underestimate the true strength of these relationships. Future studies integrating multiscale imaging and molecular data within the same cohort will be important for validating and refining these relationships. Finally, considering the well-known influence of analytic choices on functional network construction [73–77], future research should examine the generalizability of the present findings across alternative preprocessing strategies, parcellation schemes, and connectivity definitions. Such systematic validation will be essential for establishing the robustness and broader applicability of the FC allometry reported here.

## Data availability

All data that support the findings of this study are from publicly available datasets (HCP-YA resting-state and task datasets: https://www.humanconnectome.org/; SLEM dataset: http://fcon_1000.projects.nitrc.org/indi/retro/southwestuni_qiu_index.html; Neurotransmitter receptor and transporter maps: https://github.com/netneurolab/neuromaps; Gene expression data: http://human.brain-map.org/; Metabolism-related maps: https://github.com/netneurolab/neuromaps; Meta-analytic activation maps: https://github.com/neurosynth/neurosynth; ABIDE dataset: https://fcon_1000.projects.nitrc.org/indi/abide/).

## Code availability

All codes used to process resting-state fMRI data are from publicly available toolboxes, including the SPM12 toolbox (https://www.fil.ion.ucl.ac.uk/spm/software/spm12) and GRETNA toolbox (https://www.nitrc.org/projects/gretna/). Codes for all analyses of FC allometry are available at https://github.com/Changwen-Wu/Allometric_Scaling.

## Acknowledgements

This work was supported by Brain Science and Brain-like Intelligence Technology - National Science and Technology Major project (2021ZD0200500), the National Natural Science Foundation of China (Nos. 82472092 and 82502452), and grant from Research Center for Brain Cognition and Human Development, Guangdong, China (No. 2024B0303390003).

## Author contributions

J Wang conceptualized and designed the study; C Wu, J Li, S Jin, and X Jiang analyzed and interpreted the data; C Wu wrote the manuscript; J Wang, J Li, and X Qiu revised the manuscript.

## Supporting information

**Figure S1.**
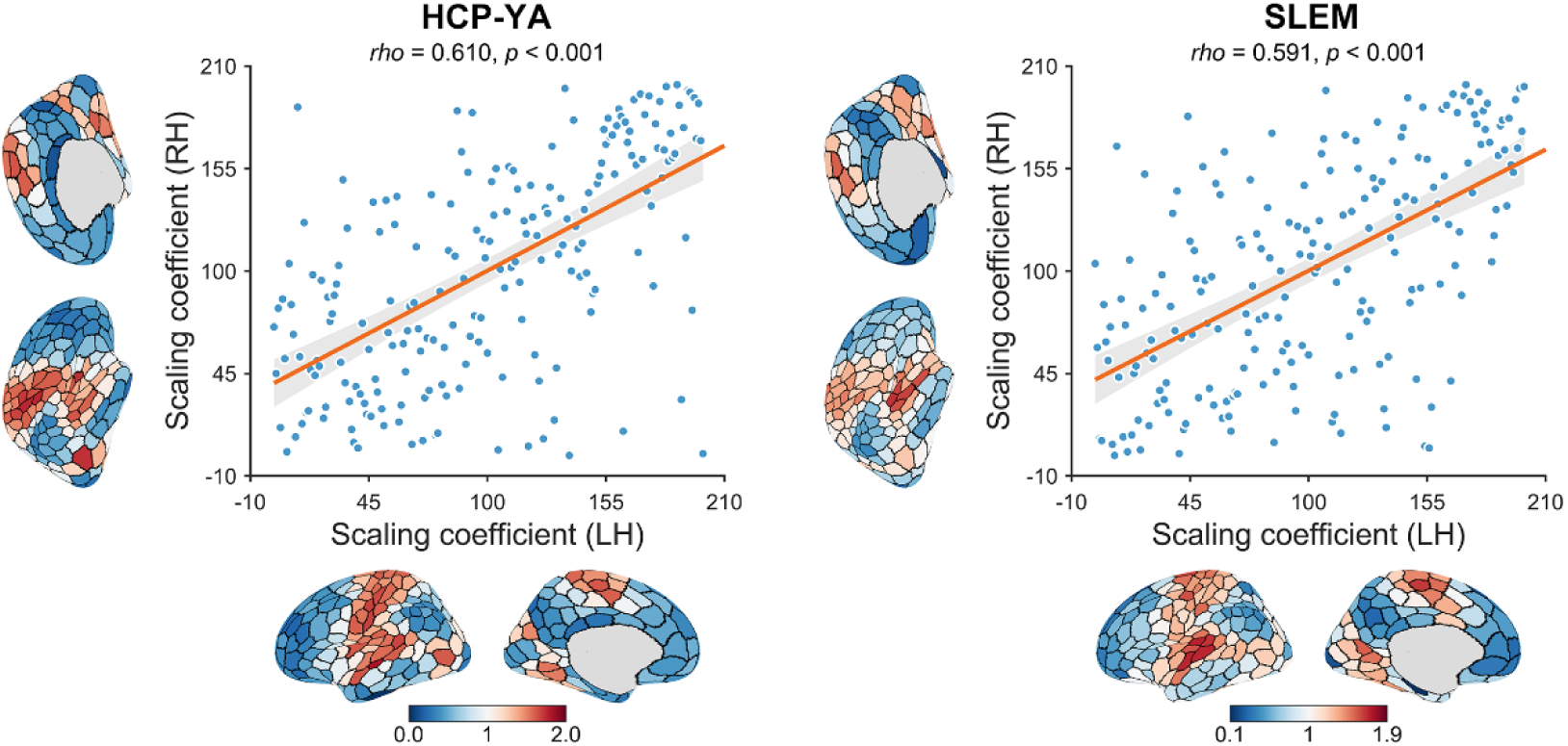
Similarity of FC allometry between hemispheres. A high positive correlation was observed in the scaling coefficients between the left and right brain hemispheres in the HCP-YA and SLEM datasets. LH, left hemisphere; RH, right hemisphere; HCP-YA, Human Connectome Project Young Adult; SLEM, Southwest University Longitudinal Imaging Multimodal.

**Figure S2.**
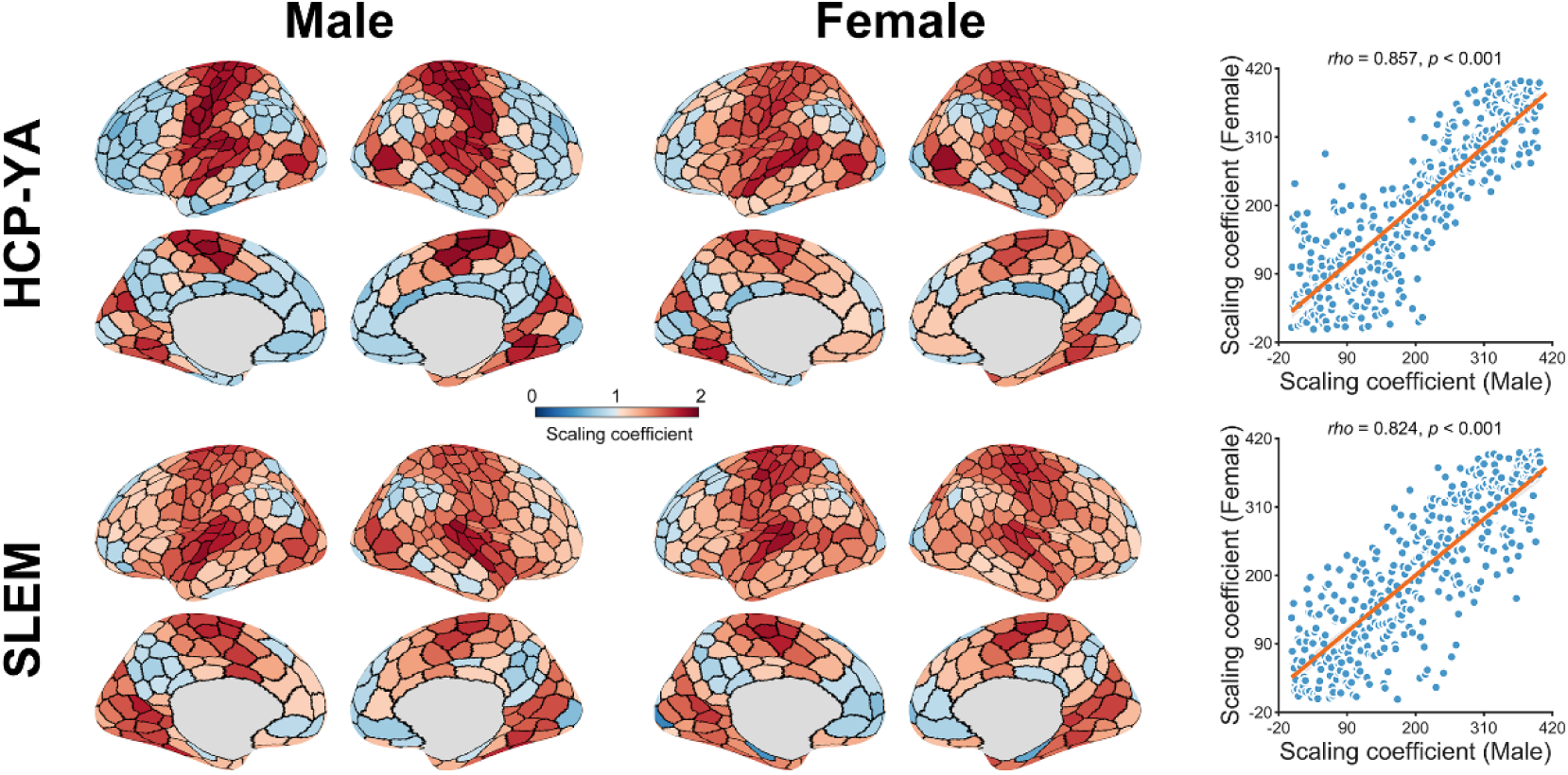
Similarity of FC allometry across sexes. A high positive correlation was observed in the scaling coefficients between males and females in the HCP-YA and SLEM datasets. HCP-YA, Human Connectome Project Young Adult; SLEM, Southwest University Longitudinal Imaging Multimodal.

**Table S1.**
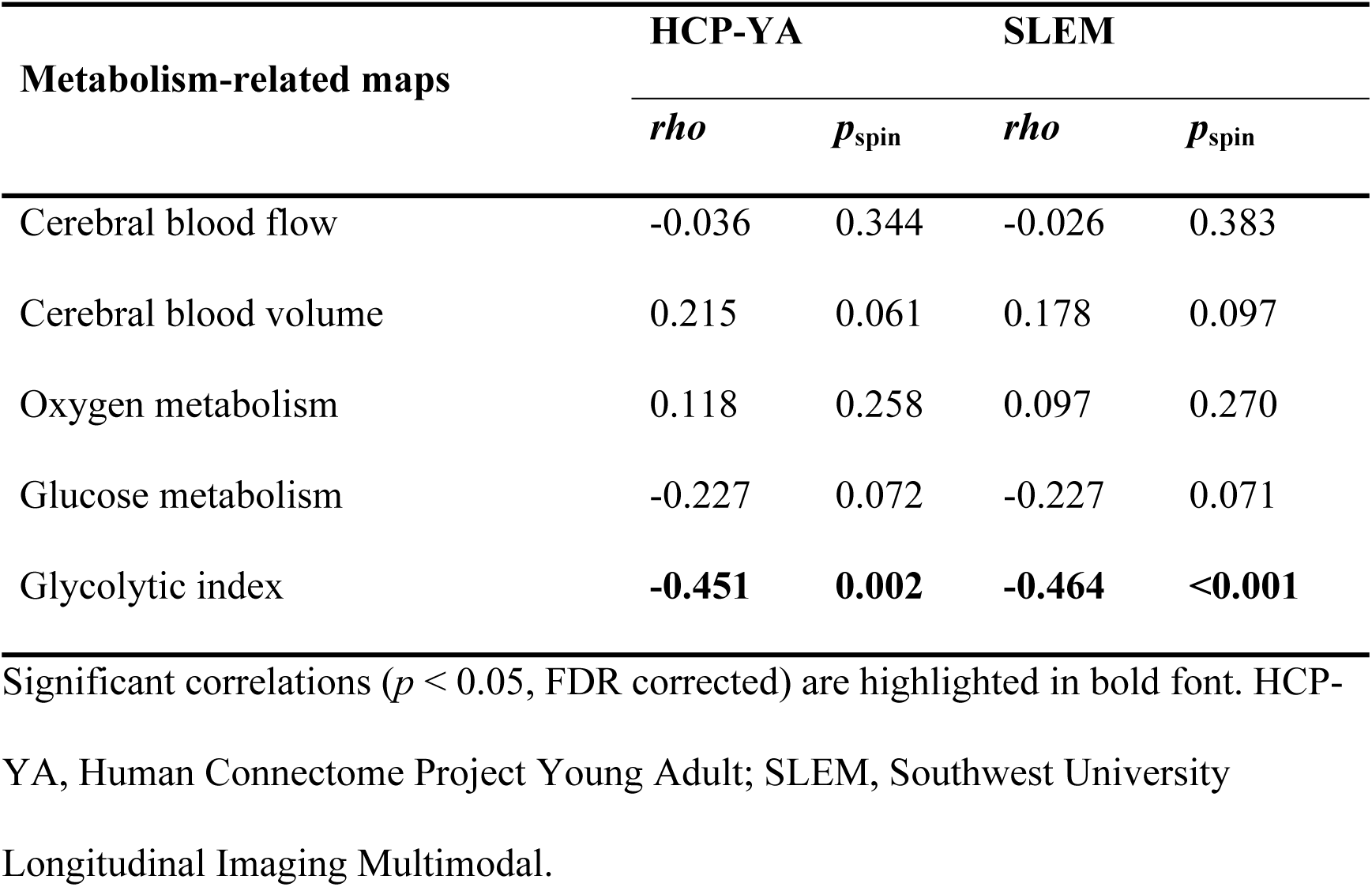
Relationship between FC allometry and metabolism-related maps.

| Metabolism-related maps | HCP-YA |  | SLEM |  |
| --- | --- | --- | --- | --- |
|  | *rho* | *p*<sub>spin</sub> | *rho* | *p*<sub>spin</sub> |
| Cerebral blood flow | -0.036 | 0.344 | -0.026 | 0.383 |
| Cerebral blood volume | 0.215 | 0.061 | 0.178 | 0.097 |
| Oxygen metabolism | 0.118 | 0.258 | 0.097 | 0.270 |
| Glucose metabolism | -0.227 | 0.072 | -0.227 | 0.071 |
| Glycolytic index | **-0.451** | **0.002** | **-0.464** | **<0.001** |

Significant correlations ( $p < 0.05$ , FDR corrected) are highlighted in bold font. HCP-
YA, Human Connectome Project Young Adult; SLEM, Southwest University
Longitudinal Imaging Multimodal.

**Table S2.** Relationship between FC allometry and neurotransmitter maps.

| Neurotransmitter maps | HCP-YA |  | SLEM |  |
| --- | --- | --- | --- | --- |
|  | *rho* | *p*<sub>spin</sub> | *rho* | *p*<sub>spin</sub> |
| Serotonin receptor 1a | -0.167 | 0.210 | -0.136 | 0.206 |
| Serotonin receptor 1b | -0.210 | 0.082 | -0.171 | 0.128 |
| Serotonin receptor 2a | **-0.428** | **<0.001** | **-0.436** | **<0.001** |
| Serotonin receptor 4 | -0.272 | 0.043 | **-0.315** | **0.013** |
| Serotonin receptor 6 | -0.068 | 0.310 | -0.051 | 0.339 |
| Serotonin transporter | 0.247 | 0.056 | 0.325 | 0.018 |
| Alpha-4 beta-2 nicotinic receptor | -0.052 | 0.367 | -0.017 | 0.437 |
| Cannabinoid receptor 1 | -0.239 | 0.066 | -0.231 | 0.037 |
| Dopamine receptor 1 | -0.171 | 0.166 | -0.093 | 0.291 |
| Dopamine receptor 2 | -0.041 | 0.434 | -0.014 | 0.476 |
| Dopamine transporter | 0.044 | 0.370 | 0.164 | 0.125 |
| γ-Aminobutyric acid type A | -0.021 | 0.478 | -0.022 | 0.453 |
| Histamine H3 receptor | -0.069 | 0.352 | -0.025 | 0.437 |
| Muscarinic 1 | -0.179 | 0.057 | -0.168 | 0.069 |
| Metabolic glutamate receptor 5 | -0.095 | 0.269 | -0.123 | 0.215 |
| Opioid receptor $\mu$ | -0.353 | 0.017 | **-0.319** | **0.011** |
| Noradrenaline transporter | **0.512** | **<0.001** | **0.500** | **<0.001** |
| Vesicular acetylcholine transporter | **0.341** | **0.008** | **0.395** | **<0.001** |

Significant correlations ( $p < 0.05$ , FDR corrected) are highlighted in bold font. HCP-
YA, Human Connectome Project Young Adult; SLEM, Southwest University
Longitudinal Imaging Multimodal.

**Table S3.** Relationship between FC allometry and gene maps.

| **Gene maps** | **HCP-YA** |  | **SLEM** |  |
| --- | --- | --- | --- | --- |
|  | *rho* | *p*<sub>spin</sub> | *rho* | *p*<sub>spin</sub> |
| Astrocytes-enriched | **-0.332** | **0.005** | **-0.355** | **0.001** |
| Endothelial cells-enriched | 0.043 | 0.419 | 0.080 | 0.280 |
| Microglia-enriched | 0.045 | 0.386 | 0.107 | 0.219 |
| Excitatory neurons-enriched | -0.206 | 0.040 | **-0.274** | **0.007** |
| Inhibitory neurons-enriched | -0.195 | 0.033 | **-0.266** | **0.004** |
| Oligodendrocytes-enriched | **0.277** | **0.001** | **0.329** | **<0.001** |
| Oligodendrocyte progenitor cells-enriched | -0.244 | 0.053 | -0.178 | 0.088 |
| Adolescence-enriched | -0.038 | 0.303 | -0.105 | 0.151 |
| Early childhood-enriched | **-0.285** | **0.003** | **-0.289** | **0.001** |
| Early fetal-enriched | 0.076 | 0.146 | 0.058 | 0.187 |
| Early middle fetal-enriched | -0.102 | 0.168 | -0.106 | 0.141 |
| Late fetal-enriched | -0.237 | 0.055 | -0.205 | 0.056 |
| Late infancy-enriched | -0.183 | 0.113 | -0.165 | 0.100 |
| Late middle fetal-enriched | -0.249 | 0.041 | -0.220 | 0.039 |
| Middle late childhood-enriched | -0.166 | 0.045 | **-0.209** | **0.017** |
| Neonatal early infancy-enriched | **-0.308** | **0.002** | **-0.308** | **0.001** |
| Young adulthood-enriched | 0.066 | 0.374 | -0.002 | 0.472 |
| Myelination-enriched | **0.228** | **0.001** | **0.207** | **0.001** |
| Axon development-enriched | **0.286** | **<0.001** | **0.279** | **<0.001** |
| Dendrite development-enriched | -0.096 | 0.125 | -0.137 | 0.054 |
| Synapse development-enriched | -0.124 | 0.026 | **-0.171** | **0.009** |

Significant correlations ( $p < 0.05$ , FDR corrected) are highlighted in bold font. HCP-
YA, Human Connectome Project Young Adult; SLEM, Southwest University
Longitudinal Imaging Multimodal.

**Table S4.** Neurosynth cognitive terms used for functional decoding of FC allometry.

|  |  |  |  |  |
| --- | --- | --- | --- | --- |
| memory | retrieval | memory retrieval | episodic memory | reasoning |
| autobiographical memory | recall | judgment | decision | cognitive control |
| thought | semantic memory | strategy | familiarity | risk |
| encoding | decision making | recognition | mood | salience |
| knowledge | impulsivity | social cognition | emotion regulation | emotion |
| context | intention | rule | uncertainty | valence |
| psychosis | intelligence | distraction | working memory | belief |
| navigation | anxiety | updating | reinforcement learning | addiction |
| priming | maintenance | extinction | task difficulty | concept |
| expectancy | goal | monitoring | search | association |
| retention | hyperactivity | stress | sentence comprehension | efficiency |
| face recognition | fear | response inhibition | eating | inference |
| reward anticipation | learning | competition | arousal | insight |
| consolidation | meaning | interference | verbal fluency | language comprehension |
| word recognition | consciousness | utility | attention | sustained attention |
| reading | manipulation | strength | anticipation | language |
| inhibition | empathy | loss | focus | sleep |
| mental imagery | visual attention | categorization | expertise | object recognition |
| effort | fixation | adaptation | communication | selective attention |
| morphology | spatial attention | response selection | balance | naming |
| rehearsal | facial expression | visual perception | induction | gaze |
| discrimination | skill | detection | listening | planning |
| pain | speech perception | integration | action | localization |
| speech production | imagery | perception | multisensory | coordination |
| rhythm | motor control | movement |  |  |

**Table S5.** Relationship between FC allometry and meta-analytic activation maps.

| **Cognitive terms** | **HCP-YA** |  | **SLEM** |  |
| --- | --- | --- | --- | --- |
|  | *rho* | *p*<sub>spin</sub> | *rho* | *p*<sub>spin</sub> |
| Memory retrieval | **-0.670** | **<0.001** | **-0.632** | **<0.001** |
| Memory | **-0.610** | **<0.001** | **-0.548** | **<0.001** |
| Retrieval | **-0.595** | **<0.001** | **-0.526** | **<0.001** |
| Cognitive control | **-0.585** | **<0.001** | **-0.530** | **<0.001** |
| Reasoning | **-0.555** | **<0.001** | **-0.560** | **<0.001** |
| Judgment | **-0.532** | **<0.001** | **-0.480** | **<0.001** |
| Episodic memory | **-0.509** | **<0.001** | **-0.474** | **<0.001** |
| Strategy | **-0.497** | **<0.001** | **-0.454** | **<0.001** |
| Autobiographical memory | **-0.494** | **<0.001** | **-0.485** | **<0.001** |
| Decision | **-0.476** | **<0.001** | **-0.442** | **<0.001** |
| Decision making | **-0.456** | **<0.001** | **-0.472** | **<0.001** |
| Rule | **-0.441** | **<0.001** | **-0.423** | **<0.001** |
| Recall | **-0.435** | **<0.001** | **-0.345** | **<0.001** |
| Thought | **-0.430** | **<0.001** | **-0.390** | **<0.001** |
| Risk | **-0.425** | **<0.001** | **-0.424** | **<0.001** |
| Impulsivity | **-0.424** | **<0.001** | **-0.414** | **<0.001** |
| Intention | **-0.404** | **<0.001** | **-0.498** | **<0.001** |
| Social cognition | **-0.380** | **<0.001** | **-0.404** | **<0.001** |
| Uncertainty | **-0.377** | **<0.001** | **-0.433** | **<0.001** |
| Knowledge | **-0.374** | **<0.001** | **-0.338** | **<0.001** |
| Salience | **-0.371** | **<0.001** | **-0.407** | **<0.001** |
| Semantic memory | **-0.360** | **<0.001** | **-0.280** | **0.001** |
| Recognition | **-0.350** | **0.002** | **-0.286** | **0.004** |
| Mood | **-0.344** | **0.001** | **-0.289** | **0.001** |
| Familiarity | **-0.339** | **<0.001** | **-0.314** | **<0.001** |
| Belief | **-0.329** | **<0.001** | **-0.385** | **<0.001** |
| Context | **-0.320** | **<0.001** | **-0.236** | **0.003** |
| Expectancy | **-0.320** | **<0.001** | **-0.304** | **<0.001** |
| Encoding | **-0.308** | **0.003** | **-0.255** | **0.010** |
| Working memory | **-0.308** | **0.004** | **-0.274** | **0.015** |
| Maintenance | **-0.296** | **0.002** | **-0.253** | **0.012** |
| Emotion regulation | **-0.293** | **0.004** | **-0.279** | **0.002** |
| Goal | **-0.268** | **0.021** | **-0.354** | **<0.001** |
| Updating | **-0.266** | **0.003** | **-0.251** | **0.004** |
| Monitoring | **-0.256** | **0.005** | **-0.231** | **0.008** |
| Emotion | -0.246 | 0.080 | -0.235 | 0.045 |
| Intelligence | **-0.244** | **0.003** | **-0.293** | **0.001** |
| Anxiety | -0.237 | 0.049 | -0.231 | 0.031 |
| Concept | **-0.231** | **0.001** | **-0.240** | **<0.001** |
| Sentence comprehension | **-0.230** | **0.005** | **-0.220** | **0.007** |
| Valence | -0.220 | 0.105 | -0.190 | 0.082 |
| Response inhibition | **-0.219** | **0.007** | -0.167 | 0.039 |
| Distraction | **-0.212** | **0.002** | -0.133 | 0.033 |
| Addiction | -0.209 | 0.051 | **-0.208** | **0.017** |
| Reinforcement learning | **-0.200** | **0.013** | **-0.209** | **0.009** |
| Efficiency | **-0.200** | **0.018** | **-0.223** | **0.010** |
| Task difficulty | -0.193 | 0.026 | -0.168 | 0.051 |
| Face recognition | -0.190 | 0.084 | -0.146 | 0.121 |
| Psychosis | -0.175 | 0.071 | -0.170 | 0.057 |
| Hyperactivity | -0.166 | 0.050 | -0.142 | 0.075 |
| Language comprehension | -0.156 | 0.043 | -0.079 | 0.179 |
| Extinction | -0.153 | 0.119 | -0.102 | 0.207 |
| Priming | -0.150 | 0.096 | -0.092 | 0.196 |
| Reward anticipation | -0.122 | 0.190 | -0.085 | 0.237 |
| Interference | -0.119 | 0.147 | -0.060 | 0.288 |
| Search | -0.116 | 0.140 | -0.118 | 0.137 |
| Meaning | -0.110 | 0.180 | -0.024 | 0.397 |
| Verbal fluency | -0.102 | 0.149 | -0.066 | 0.247 |
| Fear | -0.098 | 0.325 | -0.060 | 0.383 |
| Stress | -0.096 | 0.322 | -0.087 | 0.286 |
| Language | -0.086 | 0.290 | 0.012 | 0.475 |
| Competition | -0.076 | 0.203 | -0.081 | 0.189 |
| Eating | -0.075 | 0.366 | -0.052 | 0.373 |
| Navigation | -0.073 | 0.319 | -0.090 | 0.259 |
| Arousal | -0.072 | 0.327 | -0.082 | 0.265 |
| Retention | -0.068 | 0.192 | -0.087 | 0.130 |
| Inhibition | -0.068 | 0.271 | -0.034 | 0.376 |
| Attention | -0.065 | 0.321 | -0.095 | 0.238 |
| Association | -0.061 | 0.342 | -0.032 | 0.394 |
| Sustained attention | -0.053 | 0.275 | -0.019 | 0.408 |
| Consciousness | -0.041 | 0.308 | -0.073 | 0.182 |
| Focus | -0.033 | 0.297 | -0.063 | 0.141 |
| Empathy | -0.030 | 0.375 | -0.101 | 0.173 |
| Utility | -0.025 | 0.398 | -0.091 | 0.130 |
| Word recognition | -0.023 | 0.428 | 0.004 | 0.497 |
| Learning | -0.006 | 0.494 | 0.046 | 0.298 |
| Insight | -0.003 | 0.499 | -0.062 | 0.277 |
| Inference | -0.002 | 0.488 | -0.043 | 0.259 |
| Manipulation | 0.012 | 0.486 | 0.027 | 0.445 |
| Communication | 0.021 | 0.430 | -0.067 | 0.256 |
| Anticipation | 0.023 | 0.425 | 0.015 | 0.420 |
| Effort | 0.035 | 0.416 | 0.044 | 0.358 |
| Mental imagery | 0.036 | 0.426 | 0.079 | 0.249 |
| Expertise | 0.040 | 0.335 | 0.067 | 0.195 |
| Sleep | 0.042 | 0.321 | 0.043 | 0.309 |
| Strength | 0.044 | 0.320 | -0.010 | 0.465 |
| Loss | 0.053 | 0.315 | 0.049 | 0.306 |
| Reading | 0.055 | 0.337 | 0.121 | 0.157 |
| Response selection | 0.062 | 0.374 | 0.086 | 0.235 |
| Balance | 0.076 | 0.248 | 0.040 | 0.366 |
| Spatial attention | 0.080 | 0.349 | 0.063 | 0.333 |
| Consolidation | 0.088 | 0.185 | 0.028 | 0.361 |
| Fixation | 0.089 | 0.295 | 0.104 | 0.225 |
| Visual attention | 0.105 | 0.292 | 0.037 | 0.429 |
| Selective attention | 0.110 | 0.165 | 0.111 | 0.126 |
| Categorization | 0.139 | 0.070 | **0.219** | **0.007** |
| Rehearsal | 0.150 | 0.129 | 0.177 | 0.057 |
| Morphology | 0.158 | 0.083 | 0.211 | 0.026 |
| Listening | 0.159 | 0.177 | 0.182 | 0.110 |
| Object recognition | 0.160 | 0.172 | 0.128 | 0.179 |
| Adaptation | 0.166 | 0.077 | 0.168 | 0.048 |
| Detection | 0.172 | 0.117 | 0.150 | 0.098 |
| Induction | **0.199** | **0.020** | **0.240** | **0.001** |
| Pain | 0.205 | 0.104 | 0.181 | 0.112 |
| Planning | 0.219 | 0.107 | 0.195 | 0.077 |
| Facial expression | **0.225** | **0.017** | **0.243** | **0.002** |
| Naming | **0.237** | **0.002** | **0.257** | **0.001** |
| Skill | **0.251** | **0.011** | **0.277** | **0.001** |
| Discrimination | **0.260** | **0.006** | **0.248** | **0.002** |
| Action | 0.263 | 0.053 | 0.206 | 0.075 |
| Speech perception | **0.278** | **0.009** | **0.298** | **0.002** |
| Gaze | **0.321** | **0.006** | **0.325** | **<0.001** |
| Speech production | **0.331** | **0.004** | **0.382** | **0.001** |
| Visual perception | **0.347** | **<0.001** | **0.309** | **<0.001** |
| Localization | **0.358** | **<0.001** | **0.359** | **<0.001** |
| Integration | **0.363** | **0.002** | **0.347** | **0.001** |
| Imagery | **0.409** | **<0.001** | **0.377** | **<0.001** |
| Coordination | **0.487** | **<0.001** | **0.449** | **<0.001** |
| Motor control | **0.491** | **<0.001** | **0.458** | **<0.001** |
| Rhythm | **0.557** | **<0.001** | **0.505** | **<0.001** |
| Perception | **0.590** | **<0.001** | **0.571** | **<0.001** |
| Movement | **0.592** | **<0.001** | **0.542** | **<0.001** |
| Multisensory | **0.594** | **<0.001** | **0.547** | **<0.001** |

Significant correlations ( $p < 0.05$ , FDR corrected) are highlighted in bold font. HCP-
YA, Human Connectome Project Young Adult; SLEM, Southwest University
Longitudinal Imaging Multimodal.

**Table S6.** Demographic and clinical characteristics of ABIDE participant.

|  | **TDCs (n = 518)** | **ASD (n = 338)** | ***p*** |
| --- | --- | --- | --- |
| Age | 12.00 ± 2.79 | 12.16 ± 2.84 | 0.400<sup>a</sup> |
| Sex (F/M) | 122/396 | 66/272 | 0.164<sup>b</sup> |
| Mean framewise displacement | 0.18 ± 0.09 | 0.19 ± 0.09 | 0.200<sup>a</sup> |
| Full-scale IQ<sup>†</sup> | 111.40 ± 12.50 | 110.10 ± 14.92 | 0.191<sup>a</sup> |
| Handedness (L/Mixed/R/Missing) | 36/75/404/3 | 27/55/250/6 | 0.565<sup>b</sup> |

Data are presented as mean ± standard deviation or counts. ASD, autism spectrum disorder; TDCs, typically developing controls. F, female; M, male; IQ, intelligence quotient; L, left; R, right.
<sup>a</sup>*P*-values are obtained with two-sample *t*-tests.
<sup>b</sup>*P*-values are obtained with Chi-square tests.
<sup>†</sup>Data are available for 478 TDCs and 304 patients with ASD.

